# Sign epistasis extends the effects of balancing selection on genetic diversity

**DOI:** 10.1101/2025.04.09.647826

**Authors:** Ksenia A. Khudiakova, Nicholas H. Barton, Goran Arnqvist

## Abstract

Balancing selection, a form of selection that maintains genetic diversity, is difficult to detect, and the importance of balancing selection for the maintenance of genetic variation may be larger than often assumed. We model the possibility that the diversity-promoting effects of balancing selection extend to other loci that show sign epistasis with a locus under balancing selection. Rather than focusing on overdominance, as was done in previous efforts, we explore the effects of negative frequency dependence and show that this has important effects on the conditions under which the diversity-promoting effect of epistasis can occur in diploids. Our results show that not only recombination rate but also the dominance of sign epistasis are key parameters that determine the maintenance of polymorphism beyond the locus under direct balancing selection. We suggest that the effect we explore may play a significant role, especially when balancing selection acts on major effect loci.

## 1 Introduction

Efforts to intersect DNA sequencing data and population genetics theory with estimates of genetic variance suggest that the observed variability in fitness components cannot be maintained solely by mutation-selection balance (Barton & Keightley, 2002; Bonnet et al., 2022; B. Charlesworth, 2015). Genetic variation in life history traits is often sizeable (Burch et al., 2024; Mousseau & Roff, 1987) and within-population polymorphism in major-effect variants such as inversions are common (Faria et al., 2019). These observations call for a better understanding of the processes that act to maintain substantial levels of non-neutral genetic diversity. One possibility is that the genome-wide effects of balancing selection are larger than typically assumed (B. Charlesworth, 2015; Kroymann & Mitchell-Olds, 2005). Balancing selection (BLS) is selection that acts to maintain variation, including both overdominance (i.e., heterozygote advantage) and negative frequency-dependent selection (NFDS).

The conditions under which we expect balancing selection in a relatively restricted number of loci to affect genetic diversity more widely have been studied previously. For example, it is commonly recognized that balancing selection increases diversity across linked neutral sites (D. Charlesworth, 2006). Furher, functional dependencies between loci under balancing selection and other sites in the genome, such that different alleles are favored on different genetic backgrounds, called sign epistasis, may play a significant role in this context. Indeed, models of haploid systems confirm that epistasis can strengthen the diversifying effects of balancing selection (Kelly, 2000; Kelly & Wade, 2000).

Sign epistasis represents inter-locus interactions between alleles such that the fitness effect of a given allele in one locus changes from beneficial to deleterious depending on the allelic state at one or more other loci (fig. 1). Such epistatic interactions tend to generate complex fitness valleys, i.e., clusters of polygenic genotypes of low fitness, and fitness ridges or even distinct fitness peaks (Gavrilets, 2004; Saona et al., 2022; Poelwijk et al., 2011; Wolf et al., 2000). Epistasis has a central role in our understanding of incipient speciation, where evolving epistatic interactions within populations (Corbett-Detig et al., 2013; Ortiz-Barrientos et al., 2016) may result in reproductive isolation between populations through the build-up of epistatic Dobzhansky-Muller incompatibilities (Orr & Turelli, 2001). Yet, most evolutionary genetic models ignore interactions between loci (Hansen, 2013) and links between the role of epistasis in adaptation and its role in speciation, potentially connecting micro-and macroevolution, are not fully understood (Johnson, 2000; Wade, 2000). Although the contribution of physiological epistasis to additive genetic variance can be very substantial, epistasis is typically regarded as inconsequential for the short-term response to selection (Mäki-Tanila & Hill, 2014). Contrary to this view, epistasis can contribute to the long-term response to selection (Paixão & Barton, 2016; Bourg et al., 2024) and several experimental studies have unveiled an important role for interactions between loci in adaptation (Bloom et al., 2013; Good et al., 2017; Pettersson et al., 2011; Wolf et al., 2000). Further, it has long been recognized that epistatic variance can be “converted” into additive genetic variance following, e.g., population bottlenecks (Barton & Turelli, 2004), and line-cross analyses have shown that the epistatic contribution to genetic variance in life history traits is often substantial (Burch et al., 2024; Roff & Emerson, 2006). However, our understanding of whether and how epistasis affects those processes that maintain allelic diversity in interacting loci within populations is relatively restricted (Hansen, 2013; Hemani et al., 2013).

**Figure 1:**
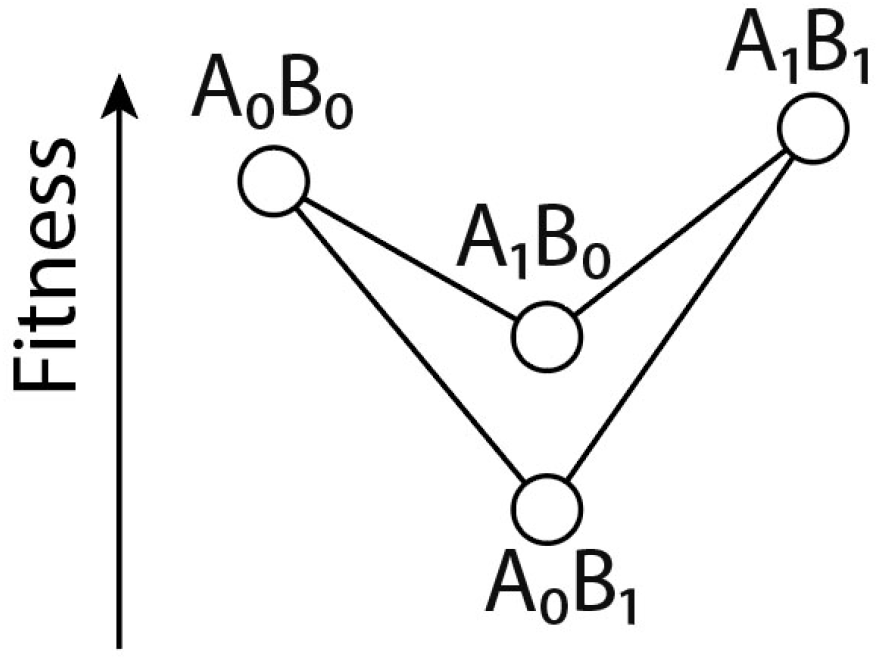
Sign epistasis between two haploid biallelic loci.

Since recombination tends to disassociate favorable allelic combinations, the potential of epistasis to maintain genetic variation depends on the balance between the strength of selection and the rate of recombination. Hence, we expect two related effects: 1) the efficacy of balancing and epistatic selection in maintaining variation should decrease with physical distance, and 2) epistatic selection will lead to evolution of tighter linkage. Already Fisher (1930, p. 103) argued that sign epistasis can lead to decreased recombination. A formal analysis was provided by Kimura (1956), who investigated the equilibria and stability of a deterministic two-locus model in which one locus exhibits heterozygote advantage and interacts with a second locus through sign epistasis. He showed that the maintenance of polymorphism at both loci is possible when linkage is sufficiently tight. Later work generalised Kimura’s fitness table in a class of fitness tables called “symmetric viability models”, i.e. fitness tables that are symmetric with respect to the fitness of the double heterozygote upon relabeling of loci and alleles (Bodmer & Felsenstein, 1967; Karlin & Feldman, 1970; Christiansen, 1999). Another inherent feature of the symmetric viability models is constant fitnesses, with balancing selection being modeled by heterozygote advantage. For all symmetric viability models, the conclusions remained similar to Kimura’s: the diversity-promoting effects of sign epistasis are restricted to the cases when loci are tightly linked, which predicts that this effect may be of limited evolutionary significance.

Here, we provide an exploration beyond symmetric viability models where we relax the assumption of constant fitnesses: instead of overdominance, we consider negative frequency dependence as the cause of balancing selection. We demonstrate that when epistatic effects show dominance, even free recombination allows for stable coexistence of polymorphic loci, challenging the long-standing view that tight linkage is required. Our results help clarify the conditions under which sign epistasis can contribute to the maintenance of within-population genetic diversity.

## 2 Materials and methods

We consider the deterministic dynamics of two biallelic loci under selection and recombination in an effectively infinite population. Locus *A* with alleles *A*_0_ and *A*_1_ is under symmetric negative frequency-dependent selection with selection coefficient *t*. Locus *B* with alleles *B*_0_ and *B*_1_ is in sign epistasis with locus *A*, which means that the genotypes containing a combination of conflicting alleles *A*_0_ and *B*_1_ or *A*_1_ and *B*_0_ experience an additional disadvantage proportional to the epistatic selection coefficient *s*. We denote the frequency of allele *A*_0_ (resp. *B*_0_) by *q*_*A*_ (*q*_*B*_) and the frequency of allele *A*_1_ (resp. *B*_1_) by *p*_*A*_ (*p*_*B*_).

For haploids, this results in the following fitness table:

| | $B_0$ | $B_1$ |
| --- | --- | --- |
| $A_0$ | $1 - tq_A$ | $1 - tq_A - s$ |
| $A_1$ | $1 - tp_A - s$ | $1 - tp_A$. |

For diploids, two additional parameters that describe the dominance relationship within the two loci are possible. Here, we assume that allelic effects at the *A* locus are additive, such that the fitness of the heterozygote *A*_0_*A*_1_ is intermediate to the fitnesses of the two homozygotes. The dominance relationship at locus *B* is arbitrary, and we model it with the parameter *α*. When *α* = 0, selection at locus *B* is additive. When *α >* 0, this generates dominance such that the fitness of each *B* heterozygote is closer to the fitter two-locus homozygote (*B*_0_ shows dominance in one background and *B*_1_ in the other). We term this dominance of the epistatic effect. Note that we consider *α < s/*2, such that there is no direct overdominance at locus *B*. Nevertheless, as we will see later, 0 *< α < s/*2 can further stabilize double polymorphism. Thus, we consider the following fitness table:

| | $B_0B_0$ | $B_1B_0$ | $B_1B_1$ |
| --- | --- | --- | --- |
| $A_0A_0$ | $1 - tq_A + s/2$ | $1 - tq_A + \alpha$ | $1 - tq_A - s/2$ |
| $A_1A_0$ | $1 - t/2$ | $1 - t/2 + \alpha$ | $1 - t/2$ |
| $A_1A_1$ | $1 - tp_A - s/2$ | $1 - tp_A + \alpha$ | $1 - tp_A + s/2$. |

We assume random mating and no sex differences in fitness. Recombination rate between two loci is denoted by *r*, which means that proportion *r* of gametes in the new generation will contain combinations of alleles different from those of the parents.

We study the dynamics analytically using the multilocus algebra introduced in Barton & Turelli (1991) and Kirkpatrick et al. (2002).

## 3 Results

### 3.1 The haploid case

The haploid dynamics are described by the following set of equations:

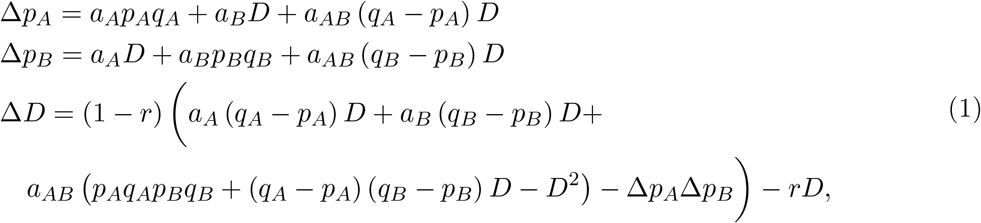

where

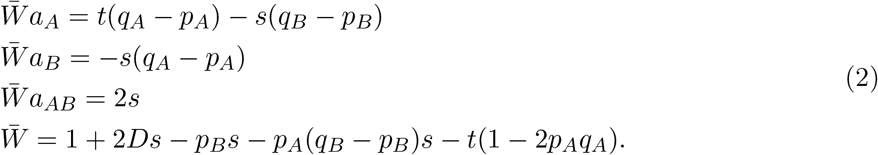

An example of the dynamics is shown on fig. 4a, SM. The system 1 admits a double-polymorphic equilibrium with *p*_*A*_ = *p*_*B*_ = 1*/*2 and the value of linkage disequilibrium

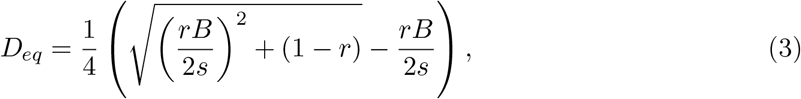

where *B* = (2 − *s* − *t*).

This equilibrium is stable when

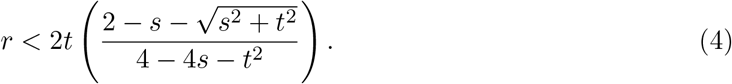

Assuming weak selection, the stability condition reduces to *r* ≲ *t*.

Numerical analyses of the selection equations confirmed these results (figs. 4b and 5, SM).

### 3.2 The diploid case

The diploid dynamics are described by the following set of equations.

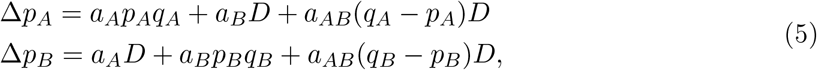

where

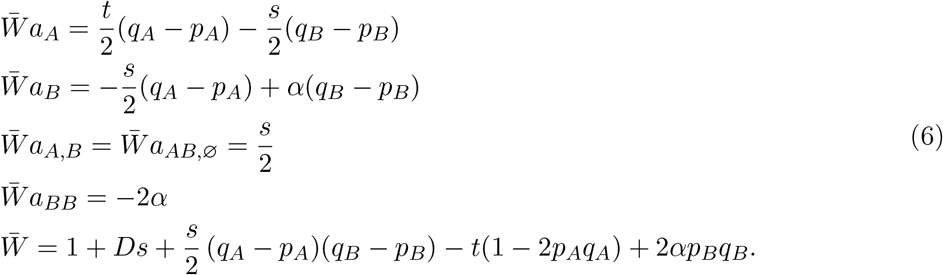

Note that in the selection coefficients, the terms involving *s* and *t* are half of those in the haploid case.

For linkage disequilibrium in diploids, we distinguish two types of statistical associations: those on the same chromosome will be denoted *D*_*AB*,ø_, and those on different chromosomes will be denoted *D*_*A,B*_. Epistatic selection builds up *D*_*A,B*_, while each round of free recombination brings *D*_*A,B*_ towards zero. The total change in *D* is given by

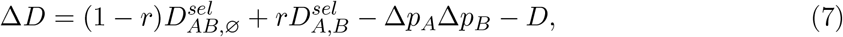

where

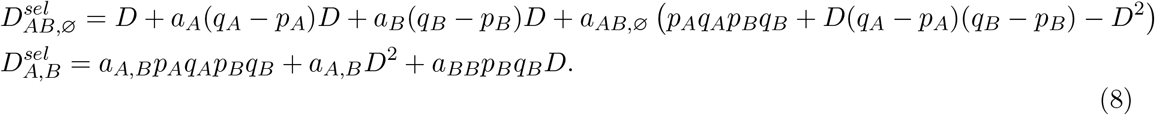

The system (5, 7) admits double-polymorphic equilibrium (*p*_*A*_ = *p*_*B*_ = 1*/*2) with

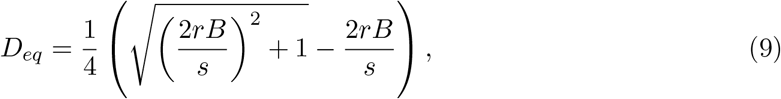

where *B* = (2 − *t* + 2*α*).

With *s < t* and *α >* 0, the double-polymorphic equilibrium is stable whenever

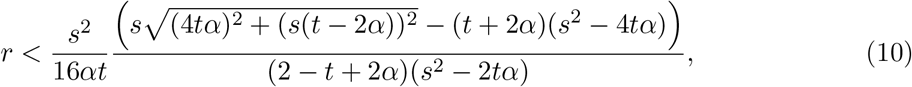

and

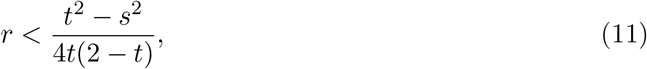

when *α* = 0. The analytical results match the numerical analysis of selection equations exactly (figs. 6 and 7, SM). As in the haploid case, thus, the condition for a double-polymorphic equilibrium in diploids under *α* = 0 requires that recombination is low relative to selection. When *s > t* such that epistatic selection is stronger than balancing selection, the double-polymorphic equilibrium is stable under condition 10 and only if *α* is large enough (fig. 2b). Increasing *α* increases the amount of recombination that can be tolerated without destabilizing the doublepolymorphic equilibrium (fig. 2).

**Figure 2:**
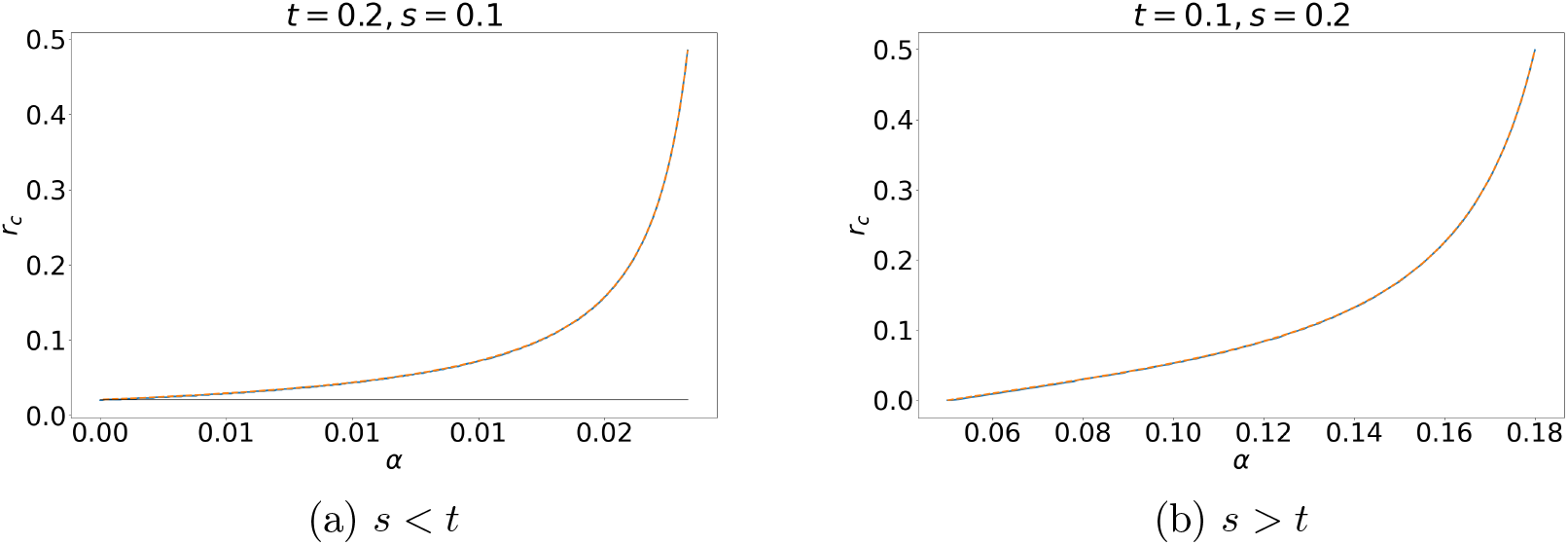
The recombination rate that can be tolerated without destabilizing the doublepolymorphic equilibrium increases with the dominance of epistatic effects *α*. The figure shows critical *r*, below which the double polymorphic equilibrium is stable, against *α* in cases where (a) balancing selection is stronger than epistatic selection and (b) epistatic selection is stronger than balancing selection. Solid line represents numerical analyses and dashed lines analytical result. When balancing selection is stronger than epistatic selection and *α* = 0, *r*_*c*_ = (*t*^2^ − *s*^2^)*/*(4*t*(2 − *t*)) (black line). When epistatic selection is stronger than balancing selection, double-polymorphic equilibrium is not stable under low values of *α*.

The increase of the parameter range with a stable double-polymorphic equilibria with an increasing *α* has an intuitive interpretation. When locus *A* is at equilibrium (*p*_*A*_ = *q*_*A*_ = 0.5), the genotypes *A*_0_*A*_0_*B*_0_*B*_0_ and *A*_1_*A*_1_*B*_1_*B*_1_ are the most-fit ones. However, due to their coexistence in a large population and random mating, less fit heterozygote *A*_1_*A*_0_ genotypes will be generated. Among those, *A*_1_*A*_0_*B*_1_*B*_0_ are the fittest whenever *α >* 0. Therefore, the reason for the stabilizing effect of *α* is essentially selection in favor of double heterozygotes that manifests itself when locus *A* is at or close to equilibrium.

## 4 Discussion

Whether sign epistasis, a form of interaction between alleles across loci where certain combinations confer high fitness, can promote genetic diversity within populations is a long-standing question. A necessary condition for this effect is the existence of stable polymorphism in a region under balancing selection. Because recombination tends to break up associations between favorable combinations of alleles (Hansen, 2013), recombination tends to decrease the stability of double-polymorphic equilibria and the general conclusion from the analysis of symmetric viability models is that linkage must be tight for double polymorphic equilibrium to be stable. Our model is not a symmetric viability model due to negative frequency-dependence. We can, nevertheless, compare our findings with previous results at equilibrium for the diploid case (the focus in previous work) by setting *p*_*A*_ = *q*_*A*_ = 1*/*2. Indeed, we recover our results for *D*_*eq*_ (eq. 9) in Kimura (1956) if we set *α* = 0, or in Karlin & Feldman (1970) and Bodmer & Felsenstein (1967) for any *α*.

As for the stability conditions, Kimura (1956), Karlin & Feldman (1970) and Bodmer & Felsenstein (1967) found that a stable coexistence of two highly adapted genotypes is only possible if linkage is tighter than balancing selection (*r* ≲ *t*). We recovered this conclusion for the haploid case, but this differs dramatically from what we observe in diploids where the parameter *α* tunes between additive *×* additive (when *α* = 0) and additive *×* dominant (when *α >* 0) effects in the fitness table. An intuitive way to illustrate the effect of *α* is to note that when *α >* 0, the fitness of the heterozygote for the *B* locus is closer to *B*_1_*B*_1_ in a *A*_1_*A*_1_ homozygote background and closer to *B*_0_*B*_0_ in a *A*_0_*A*_0_ homozygote background (fig. 3). This illustrates why we expect *α >* 0 to stabilize double polymorphism. The same stabilizing effect can also occur in general symmetric viability models (Bodmer & Felsenstein, 1967; Karlin & Feldman, 1970), where there is enough flexibility in the parameters to allow for arbitrary dominance relationships within loci. However, in symmetric viability models, the dominance of epistasis does not increase the critical recombination rate as much as in the model with frequency-dependent selection, and it seems that the critical importance of epipstatic dominance was therefore overlooked in previous contributions.

**Figure 3:**
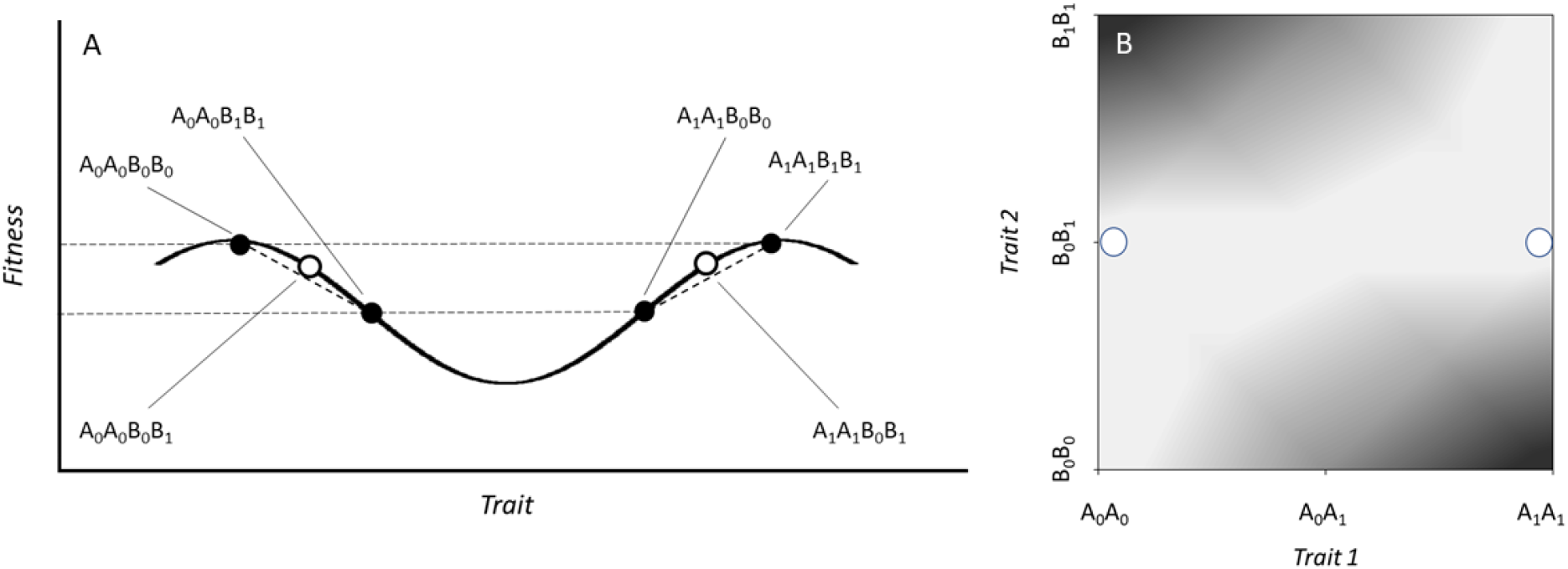
For polygenic traits under non-linear selection, epistatic interactions between loci are inevitable, and curved fitness landscapes will then lead to complex allelic effects across loci. Shown are two schematic and simplified illustrations of ways in which dominance of sign epistasis could emerge. In these, genotype determines individual trait values which in turn translate into individual fitness. Under balancing selection, the relation between fitness and trait depends on allele frequencies. To simplify illustration, the frequencies at locus *A* are here fixed at *p*_*A*_ = *q*_*A*_ = 1*/*2. (A) When two loci have additive effects on a single phenotypic trait with two distinct fitness peaks (i.e., non-linear disruptive selection), the fitness of the heterozygotes for the *B* locus (open circles) may be closer to the homozygote *B*_0_*B*_0_ in the *A*_0_*A*_0_ background and closer to *B*_1_*B*_1_ in the *A*_1_*A*_1_ background, such that the dominance of the two *B* alleles is reversed across *A* genetic backgrounds (*A*_0_*A*_1_ heterozygotes not included here, for simplicity). (B) Alternatively, for two loci coding for distinct traits, correlational selection will also result in epistasis for fitness (light - high relative fitness, dark - low relative fitness). This may also reverse the dominance of the two *B* alleles across *A* genetic backgrounds as a result of the curvature of the fitness landscape. In this example, the fitness of the heterozygotes for the *B* locus (open circles) is closer to the homozygote *B*_0_*B*_0_ in the *A*_0_*A*_0_ background and closer to *B*_1_*B*_1_ in the *A*_1_*A*_1_ background.

**Figure 4:**
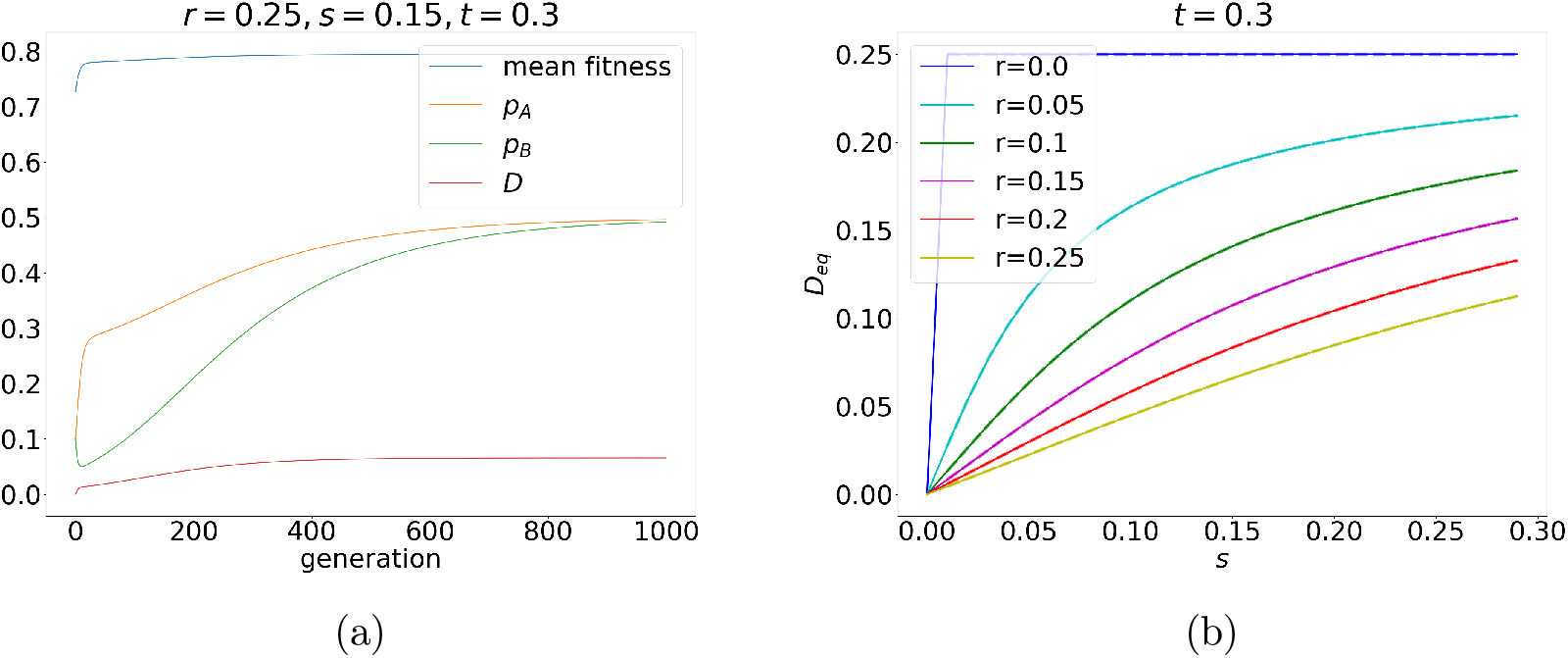
(a) Change of *p*_*A*_, *p*_*B*_, *D* and mean fitness over time and (b) the relationship between the strength of epistatic selection *s* and linkage disequilibrium *D*. Solid lines represent numerical iteration of selection equations and dashed lines analytical prediction (3).

**Figure 5:**
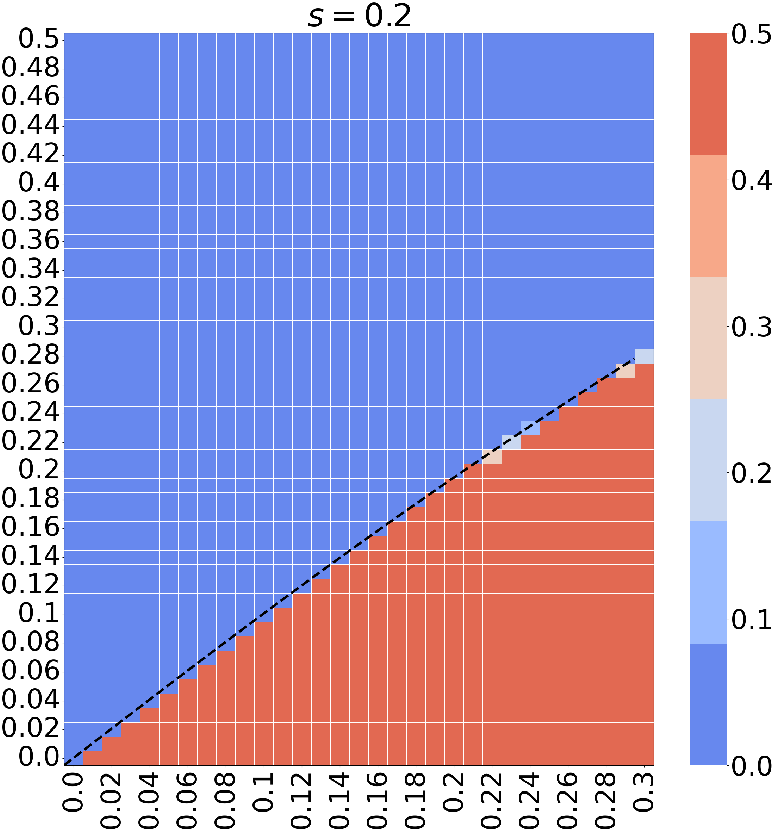
Equilibrium frequency of allele *B*_1_ in the numerical iteration of selection equations on the (*r, t*) space. Dashed line represents stability condition (4).

**Figure 6:**
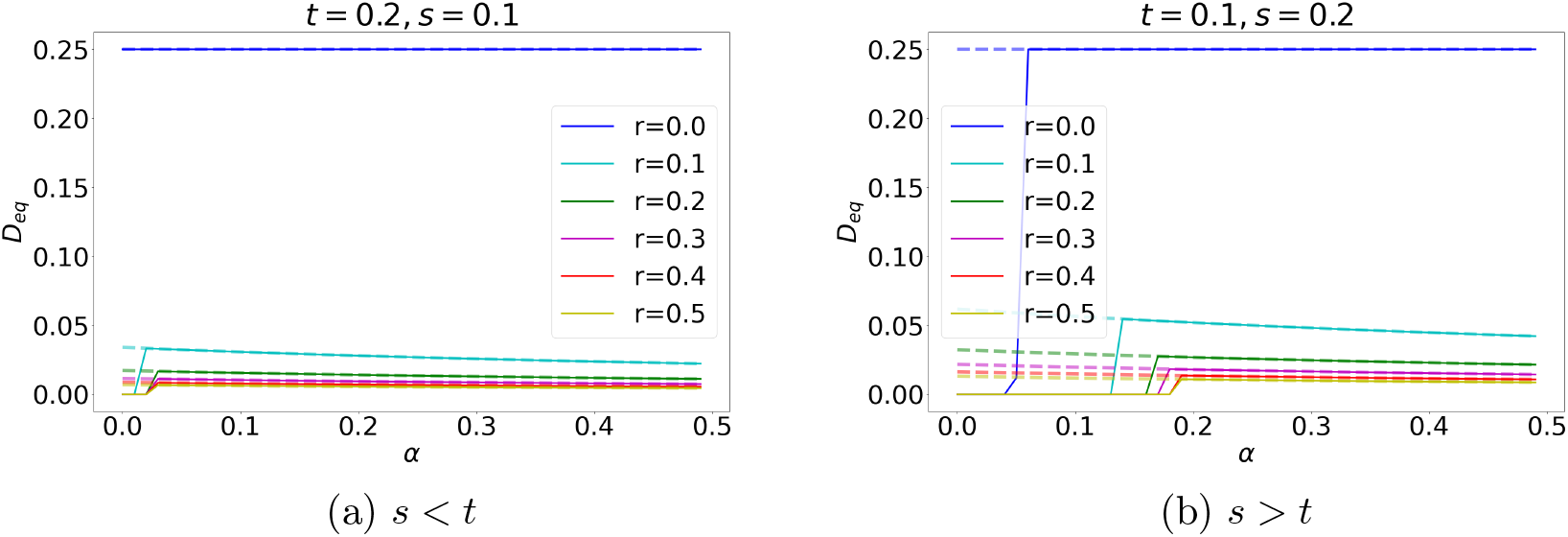
Linkage disequilibrium *D* at double-polymorphic equilibria decreases with increasing recombination rate, but is contingent upon the strength of selection and epistatic dominance *α*. Conditions where (a) balancing selection is stronger than epistatic selection and (b) epistatic selection is stronger than balancing selection. Solid lines represent numerical analysis and dashed lines are analytical prediction (9).

**Figure 7:**
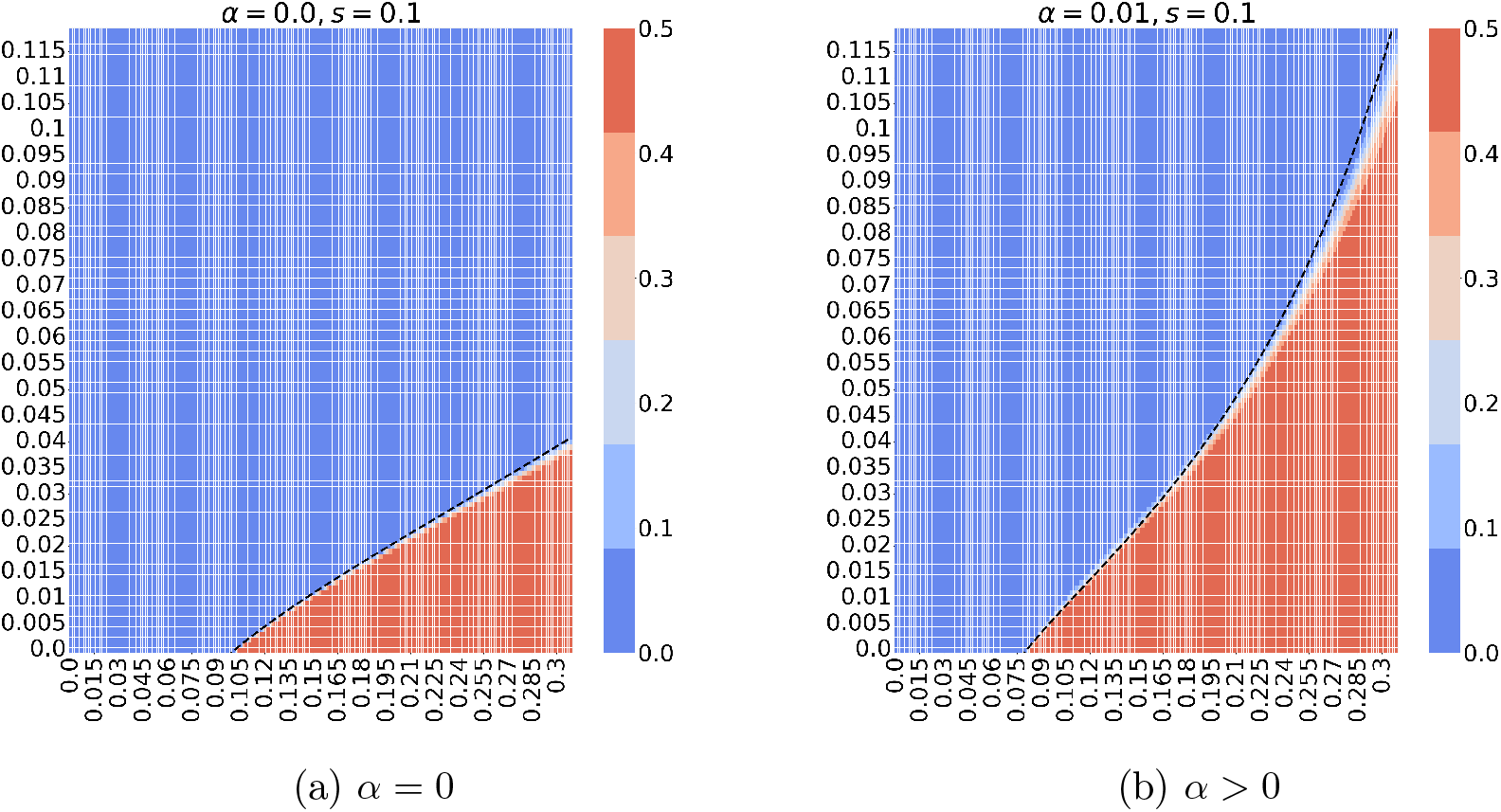
Equilibrium *p*_*B*_ in the iteration of selection equations on the (*r, t*) space. The dashed lines are critical *r* below which the double polymorphism is predicted to be stable: *r*_*c*_ = (*t*^2^ − *s*^2^)*/*(8*t* − 4*t*^2^) when *α* = 0 and as in (10) when *α >* 0.

By analogy with a general discussion of context-specific dominance effects (Connallon & Chenoweth, 2019; Curtsinger et al., 1994), we may refer to the case *α >* 0 as a beneficial reversal of dominance of epistasis because the fitness of the *B* heterozygote will be similar to the most fit homozygote across different genetic backgrounds. We note that this is in spirit analogous to dominance reversals across other contexts (e.g., environments, sexes, etc), which are known to promote stable polymorphism in single locus scenarios (Connallon & Chenoweth, 2019). Drawing on parallels in single locus scenarios (Siljestam et al., 2024; Spencer & Priest, 2016), we suggest that selection should favor dominance of sign epistasis since it effectively reduces segregation load. Because the process we describe here is predicted to extend the life span of alleles across interacting loci, it may provide sufficient time to allow the evolution of elevated dominance of epistasis which, in turn, will tend to further stabilize double-polymorphism. We note that context-specific dominance effects can emerge from a variety of non-linearities at any level of the genotype-to-phenotype map (Billiard et al., 2021).

### Sources of epistasis

The scenario studied here can be seen as representing a variety of mechanisms for the emergence of sign epistasis. This can appear on a genetic level in the form of co-adapted amino acids or nucleotide substitutions (Kondrashov et al., 2002) or more generally as co-adapted gene complexes (Dobzhansky & Pavlovsky, 1958). A general possibility is the emergence of sign epistasis for fitness as a result of non-linear fitness functions for polygenic traits (fig. 3). For example, it has been suggested that major effect life history genes may frequently experience balancing selection (Arnqvist & Rowe, 2023; Christie & McNickle, 2023) and the processes we describe may act to maintain variation in loci that show epistasis with such genes. There are many potential examples of the scenario we explore (Arnqvist & Rowe, 2023; Christie & McNickle, 2023). A common case may be inversion polymorphism maintained by balancing selection (Berdan et al., 2022), where linkage disequilibrium between the inversion and other loci are commonly observed and actually constituted a very early demonstration of epistatic selection (Prakash & Lewontin, 1968). More specific examples include a polymorphic major effect life history locus located on the sex-chromosomes in *Drosophila mercatorum*, which interacts with several autosomal modifiers all of which are highly polymorphic in natural population (Templeton, 2000). A more familiar case is color pattern in male guppies, which shows high levels of standing genetic variation maintained by negative frequency-dependent selection. New work suggests that although a major effect locus is located on the Y-chromosome, color pattern is determined by an epistatic interaction between Y and an autosomal region in LD with the Y (Paris et al., 2022). Similarly, major effect variants that affect coloration in butterfly mimicry systems maintained by balancing selection have epistatic modifiers that apparently fine-tune coloration (Nijhout, 1991), and balancing selection on inter-locus allele combinations is likely important in maintaining polymorphism in flower morphology in plants (Kelly, 2000). A distinct but classic example may in fact be sickle-cell anemia in humans, which is controlled by the major effect HBB locus well known to be under balancing selection. It is perhaps less well known that the phenotypic manifestation of HBB genotype is determined by epistatic interactions with several other polymorphic loci (Templeton, 2000). Another illustrative example comes from supergenes (chromosomal inversions) that control social behavior in some ants and are maintained as balanced polymorphisms (Purcell & Brelsford, 2025). In the ant *Formica cinerea*, a second supergene on a different chromosome influences body size and is likewise polymorphic. Despite being unlinked, the two supergenes show strong linkage disequilibrium, which appears to be maintained by strong epistatic selection (Scarparo et al., 2026). Finally, an analogue to the scenario we explore is loci under sexually antagonistic selection, where different alleles are favored in males and females (Bonduriansky & Chenoweth, 2009). Such loci essentially exhibit sign epistasis for fitness with sex-determining loci under balancing selection, which can act to maintain variation in sexually antagonistic loci given permissive patterns of dominance of epistasis (Siljestam et al., 2024).

### Possible scenarios for the accumulation of epistatic polymorphisms

Suppose that we start with locus *A* under balancing selection. Alleles at other loci favored in one genetic background but not the other may then accumulate if linkage is tight enough. This will generate indirect selection for reduced recombination rate (Otto & Feldman, 1997), a scenario analogous to the canonical model of sex chromosome evolution. Indeed, our analysis of the haploid case shows that the stability of a double polymorphism requires *r* ≲ *t*.

However, we show that in diploids, the situation is significantly different because of possible dominance effects. Given balancing selection on locus *A*, there are at least two scenarios where these sorts of effects could be important. First, in cases where epistatic balancing selection maintains two-locus polymorphism for long enough, selection will favor the evolution of “beneficial” dominance effects (i.e., selection will favor the evolution of larger *α*), with effects similar to reduced recombination. This will, in turn, further stabilize the polymorphism. Second, as new mutations with epistatic effects enter into a population, those with permissive epistatic dominance effects (i.e., *α >* 0) will tend to be retained by epistatic balancing selection. This way, selection may act as a sieve favoring the maintenance of, or increased lifespan of, mutations showing overall *α >* 0. In both cases, we would expect the build-up of genetic polymorphism and *α >* 0 to be common. Needless to say, the build-up of genetic polymorphism will be generally promoted by linkage, but as we show here, dominance effects make close linkage a less critical requirement.

### How to detect the effect in real data?

Uncovering the effects we discuss here using genome scans or population genomic data alone will clearly be very challenging (Arnqvist & Rowe, 2023). Co-adaptation of alleles at different loci should lead to positive linkage disequilibrium between them and the tools available for identifying regions under balancing selection are improving (Bitarello et al., 2023). Scanning for loci under balancing selection and inspecting regions or sites in positive LD with those may therefore be rewarding. For example, Stolyarova et al. (2022) found pervasive elevated short-range LD between nonsynonymous sites within divergent intra-genic haploblocks in the fungus *Schizophyllum communae*. This finding suggests that this may be due to a combination of balancing and epistatic selection, potentially similar to the scenario we model here. Another possibility may be to empirically estimate the dominance of epistasis (i.e., *α*), through for example advanced line-cross analyses (Burch et al., 2024), as this is expected to be positive under stable double-polymorphism. Genomic time-series data may also offer interesting routes to assess epistatic selection, as manifested by synchronous frequency shifts in populations.

In summary, we delineate the conditions under which the diversity-promoting effect of balancing selection is expected to extend to other loci through epistasis. We find that the dominance of epistasis is central in this context, and a diversifying effect is predicted even under fairly high rates of recombination if this dominance is sizeable.

## Data availability

The Python code used for the numerical analyzes is available online at https://github.com/khudyakovaks/Sign-epistasis-extends-the-effects-of-balancing-selection-on-genetic-diversity.

## Author contributions

GA, NB, and K.K. conceived and designed the analysis; NB and KK performed the analysis; GA and KK wrote the paper;

## Funding

This work was funded by grants from the Swedish Research Council (2023-03730 to G.A.) and the DOC fellowship from the Austrian Academy of Science (26293 to K.K.).

## Conflict of interest

The authors declare no conflicts of interest.

## Acknowledgments

We thank L. Rowe for valuable discussions.

## A Supplementary Material

### A.1 Haploids

### A.2 Diploids

## Notes

### Competing Interest Statement

The authors have declared no competing interest.

### Summary of Updates

The authors expanded the Introduction and Discussion to clarify how their results compare to previous works.

